# The sequence of human ACE2 is suboptimal for binding the S spike protein of SARS coronavirus 2

**DOI:** 10.1101/2020.03.16.994236

**Authors:** Erik Procko

## Abstract

The rapid and escalating spread of SARS coronavirus 2 (SARS-CoV-2) poses an immediate public health emergency. The viral spike protein S binds ACE2 on host cells to initiate molecular events that release the viral genome intracellularly. Soluble ACE2 inhibits entry of both SARS and SARS-2 coronaviruses by acting as a decoy for S binding sites, and is a candidate for therapeutic, prophylactic and diagnostic development. Using deep mutagenesis, variants of ACE2 are identified with increased binding to the receptor binding domain of S. Mutations are found across the interface, in the N90-glycosylation motif, and at buried sites where they are predicted to enhance local folding and presentation of the interaction epitope. When single substitutions are combined, large increases in binding can be achieved. The mutational landscape offers a blueprint for engineering high affinity proteins and peptides that block receptor binding sites on S to meet this unprecedented challenge.

---

In December, 2019, a novel zoonotic betacoronavirus closely related to bat coronaviruses spilled over to humans, possibly at the Huanan Seafood Market in the Chinese city of Wuhan (*1*, *2*). The virus, called SARS-CoV-2 due to its similarities with the severe acute respiratory syndrome (SARS) coronavirus responsible for a smaller outbreak nearly two decades prior (*3*, *4*), has since spread human-to-human rapidly across the world, precipitating extraordinary containment measures from governments (*5*). These events are unlike any experienced in generations. Symptoms of coronavirus disease 2019 (COVID-19) range from mild to dry cough, fever, pneumonia and death, and SARS-CoV-2 is devastating among the elderly and other vulnerable groups (*6*, *7*).

The S spike glycoprotein of SARS-CoV-2 binds angiotensin-converting enzyme 2 (ACE2) on host cells (*2*, *8*-*13*). S is a trimeric class I viral fusion protein that is proteolytically processed into S1 and S2 subunits that remain noncovalently associated in a prefusion state (*8*, *11*, *14*). Upon engagement of ACE2 by a receptor binding domain (RBD) in S1 (*15*), conformational rearrangements occur that cause S1 shedding, cleavage of S2 by host proteases, and exposure of a fusion peptide adjacent to the S2’ proteolysis site (*14*, *16*-*18*). Favorable folding of S to a post-fusion conformation is coupled to host cell/virus membrane fusion and cytosolic release of viral RNA. Atomic contacts with the RBD are restricted to the protease domain of ACE2 (*19*, *20*), and soluble ACE2 (sACE2) in which the transmembrane domain is removed is sufficient for binding S and neutralizing infection (*12*, *21*-*24*). In principle, the virus has limited potential to escape sACE2-mediated neutralization without simultaneously decreasing affinity for native ACE2 receptors, thereby attenuating virulence. Furthermore, fusion of sACE2 to the Fc region of human immunoglobulin can provide an avidity boost while recruiting immune effector functions and increasing serum stability, an especially desirable quality if intended for prophylaxis (*23*, *25*), and sACE2 has proven safe in healthy human subjects (*26*) and patients with lung disease (*27*). Recombinant sACE2 is being evaluated in a European phase II clinical trial for COVID-19 managed by Apeiron Biologics, and peptide derivatives of ACE2 are also being explored as cell entry inhibitors (*28*).

Since human ACE2 has not evolved to recognize SARS-CoV-2 S, it was hypothesized that mutations may be found that increase affinity for therapeutic and diagnostic applications. The coding sequence of full length ACE2 with an N-terminal c-myc epitope tag was diversified to create a library containing all possible single amino acid substitutions at 117 sites spanning the entire interface with S and lining the substrate-binding cavity. S binding is independent of ACE2 catalytic activity (*23*) and occurs on the outer surface of ACE2 (*19*, *20*), whereas angiotensin substrates bind within a deep cleft that houses the active site (*29*). Substitutions within the substrate-binding cleft of ACE2 therefore act as controls that are anticipated to have minimal impact on S interactions, yet may be useful for engineering out substrate affinity to enhance *in vivo* safety. However, it is important to note that catalytically active protein may have desirable effects for replenishing lost ACE2 activity in COVID-19 patients in respiratory distress (*30*, *31*).

The ACE2 library was transiently expressed in human Expi293F cells under conditions that typically yield no more than one coding variant per cell, providing a tight link between genotype and phenotype (*32*, *33*). Cells were then incubated with a subsaturating dilution of medium containing the RBD of SARS-CoV-2 fused C-terminally to superfolder GFP (sfGFP: (*34*)) (Fig. 1A). Levels of bound RBD-sfGFP correlate with surface expression levels of myc-tagged ACE2 measured by dual color flow cytometry. Compared to cells expressing wild type ACE2 (Fig. 1C), many variants in the ACE2 library fail to bind RBD, while there appeared to be a smaller number of ACE2 variants with higher binding signals (Fig. 1D). Cells expressing ACE2 variants with high or low binding to RBD were collected by fluorescence-activated cell sorting (FACS), referred to as “nCoV-S-High” and “nCoV-S-Low” sorted populations, respectively. During FACS, fluorescence signal for bound RBD-sfGFP continuously declined, requiring the collection gates to be regularly updated to ‘chase’ the relevant populations. This is consistent with RBD dissociating over hours during the experiment.

**Figure 1.**
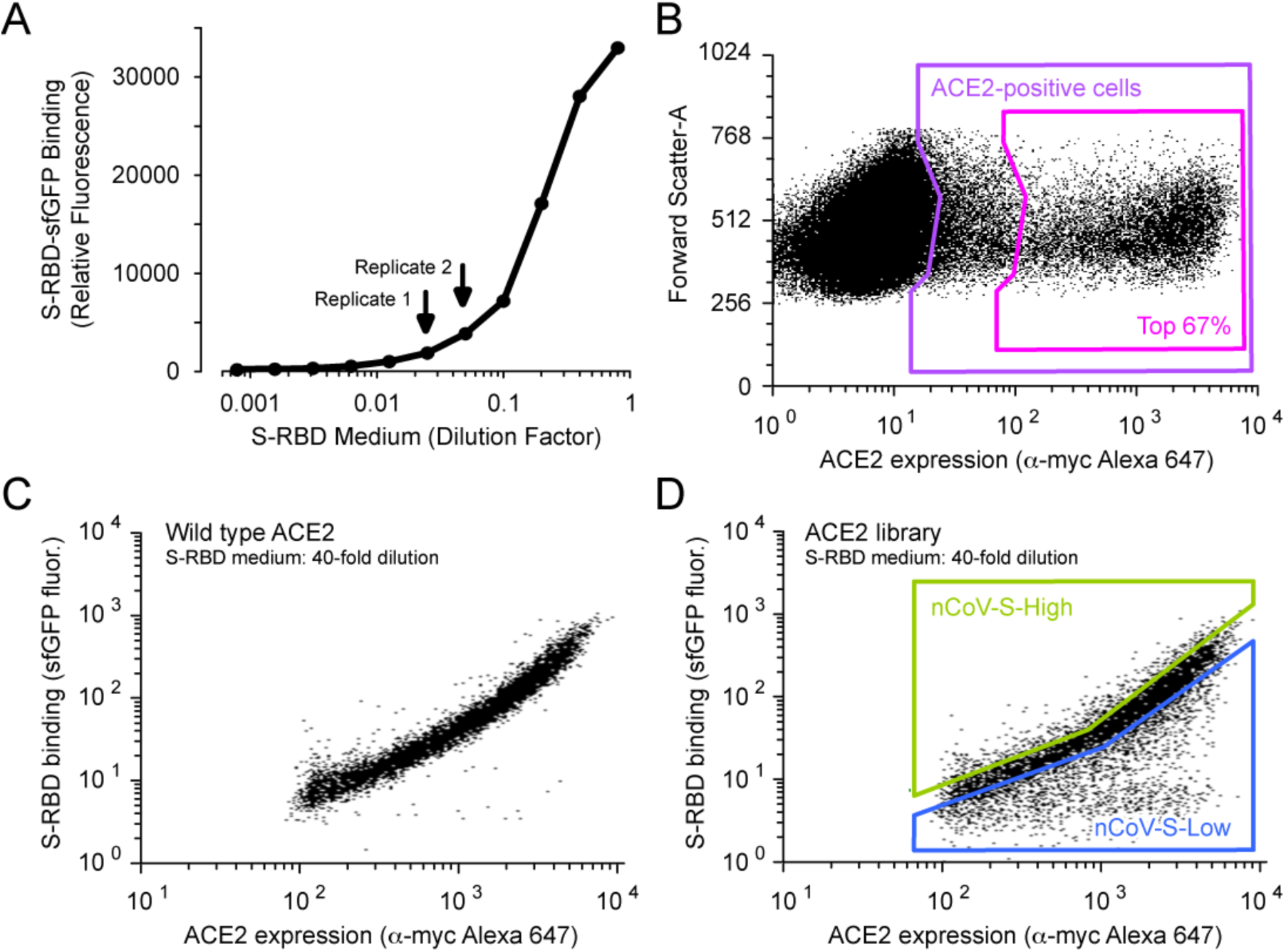
A selection strategy for ACE2 variants with high binding to the RBD of SARS-CoV-2 S. **(A)** Media from Expi293F cells secreting the SARS-CoV-2 RBD fused to sfGFP was collected and incubated at different dilutions with Expi293F cells expressing myc-tagged ACE2. Bound RBD-sfGFP was measured by flow cytometry. The dilutions of RBD-sfGFP-containing medium used for FACS selections are indicated by arrows. **(B-C)** Expi293F cells were transfected with wild type ACE2 plasmid diluted with a large excess of carrier DNA. It has been previously shown that under these conditions, cells typically acquire no more than one coding plasmid and most cells are negative. Cells were incubated with RBD-sfGFP-containing medium and co-stained with fluorescent anti-myc to detect surface ACE2 by flow cytometry. During analysis, the top 67% (magenta gate) were chosen from the ACE2-positive population (purple gate) (B). Bound RBD was subsequently measured relative to surface ACE2 expression (C). **(D)** Expi293F cells were transfected with an ACE2 single site-saturation mutagenesis library and analyzed as in B. During FACS, the top 15% of cells with bound RBD relative to ACE2 expression were collected (nCoV-S-High sort, green gate) and the bottom 20% were collected separately (nCoV-S-Low sort, blue gate).

Transcripts in the sorted populations were deep sequenced, and frequencies of variants were compared to the naive plasmid library to calculate the enrichment or depletion of all 2,340 coding mutations in the library (Fig. 2). This approach of tracking an *in vitro* selection or evolution by deep sequencing is known as deep mutagenesis (*35*). Enrichment ratios (Fig. 3A and 3B) and residue conservation scores (Fig. 3D and 3E) closely agree between two independent sort experiments, giving confidence in the data. For the most part, enrichment ratios (Fig. 3C) and conservation scores (Fig. 3F) in the nCoV-S-High sorts are anticorrelated with the nCoV-S-Low sorts, with the exception of nonsense mutations which were appropriately depleted from both gates. This indicates that most, but not all, nonsynonymous mutations in ACE2 did not eliminate surface expression. The library is biased towards solvent-exposed residues and has few substitutions of buried hydrophobics that might have bigger effects on plasma membrane trafficking (*33*).

**Figure 2.**
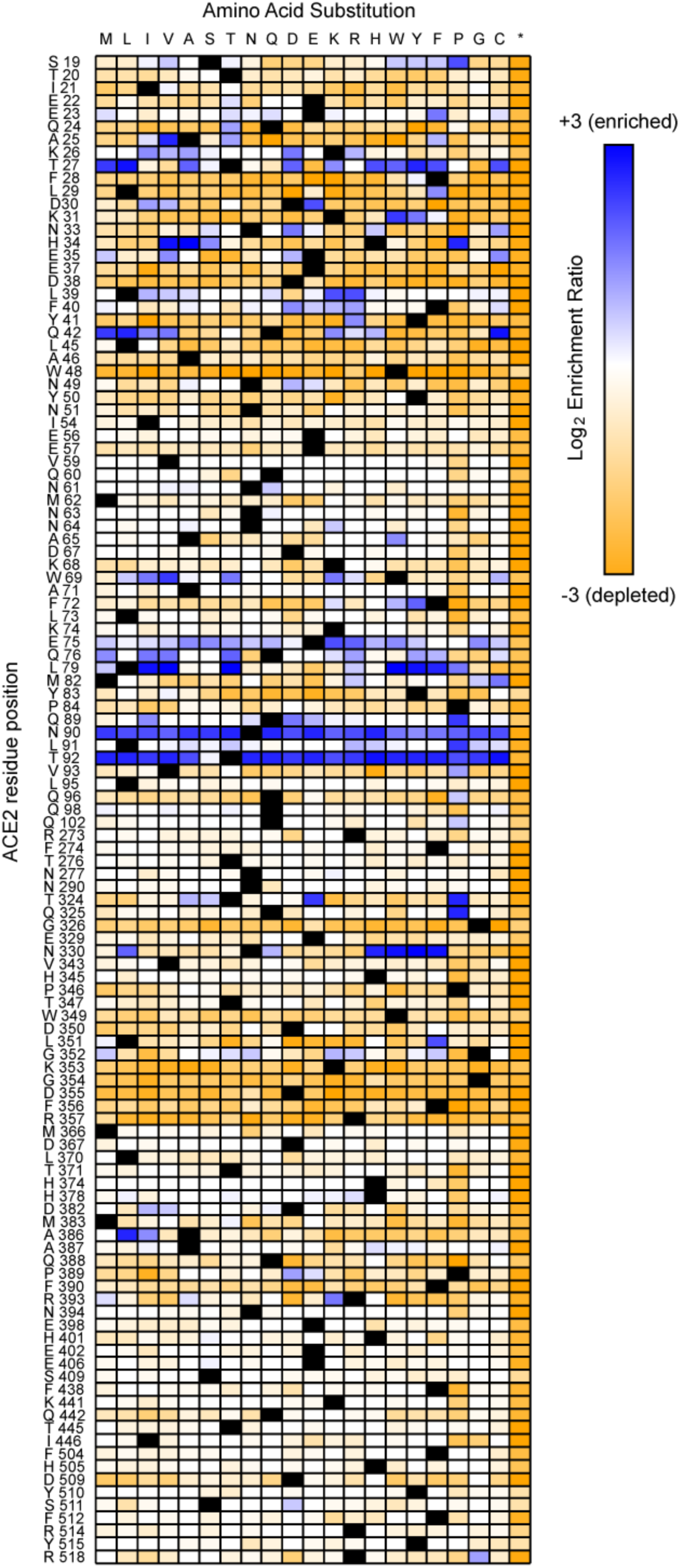
A mutational landscape of ACE2 for high binding signal to the RBD of SARS-CoV-2 S. Log_2_ enrichment ratios from the nCoV-S-High sorts are plotted from ≤ −3 (i.e. depleted/deleterious, orange) to neutral (white) to ≥ +3 (i.e. enriched, dark blue). ACE2 primary structure is on the vertical axis, amino acid substitutions are on the horizontal axis. *, stop codon.

**Figure 3.**
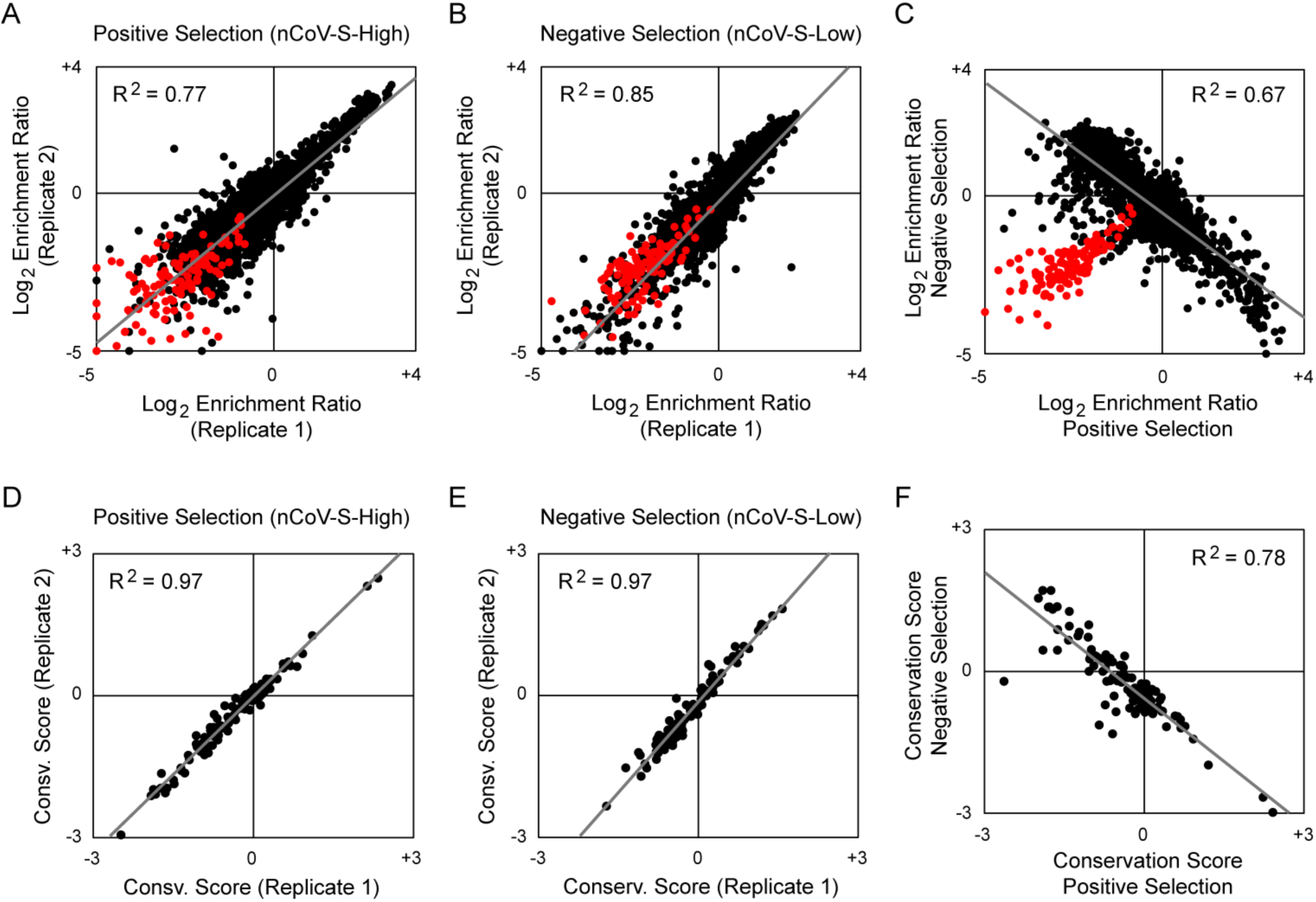
Data from independent replicates show close agreement. **(A-B)** Log_2_ enrichment ratios for ACE2 mutations in the nCoV-S-High (A) and nCoV-S-Low (B) sorts closely agree between two independent FACS experiments. Nonsynonymous mutations are black, nonsense mutations are red. Replicate 1 used a 1/40 dilution and replicate 2 used a 1/20 dilution of RBD-sfGFP-containing medium. R^2^ values are for nonsynonymous mutations. **(C)** Average log_2_ enrichment ratios tend to be anticorrelated between the nCoV-S-High and nCoV-S-Low sorts. Nonsense mutations (red) and a small number of nonsynonymous mutations (black) are not expressed at the plasma membrane and are depleted from both sort populations (i.e. fall below the diagonal). **(D-F)** Correlation plots of residue conservation scores from replicate nCoV-S-High (D) and nCoV-S-Low (E) sorts, and from the averaged data from both nCoV-S-High sorts compared to both nCoV-S-Low sorts (F). Conservation scores are calculated from the mean of the log_2_ enrichment ratios for all amino acid substitutions at each residue position.

Mapping the experimental conservation scores from the nCoV-S-High sorts to the structure of RBD-bound ACE2 (*19*) shows that residues buried in the interface tend to be conserved, whereas residues at the interface periphery or in the substrate-binding cleft are mutationally tolerant (Fig. 4A). The region of ACE2 surrounding the C-terminal end of the ACE2 α1 helix and β3-β4 strands has a weak tolerance of polar residues, while amino acids at the N-terminal end of α1 and the C-terminal end of α2 prefer hydrophobics (Fig. 4B), likely in part to preserve hydrophobic packing between α1-α2. These discrete patches contact the globular RBD fold and a long protruding loop of the RBD, respectively.

**Figure 4.**
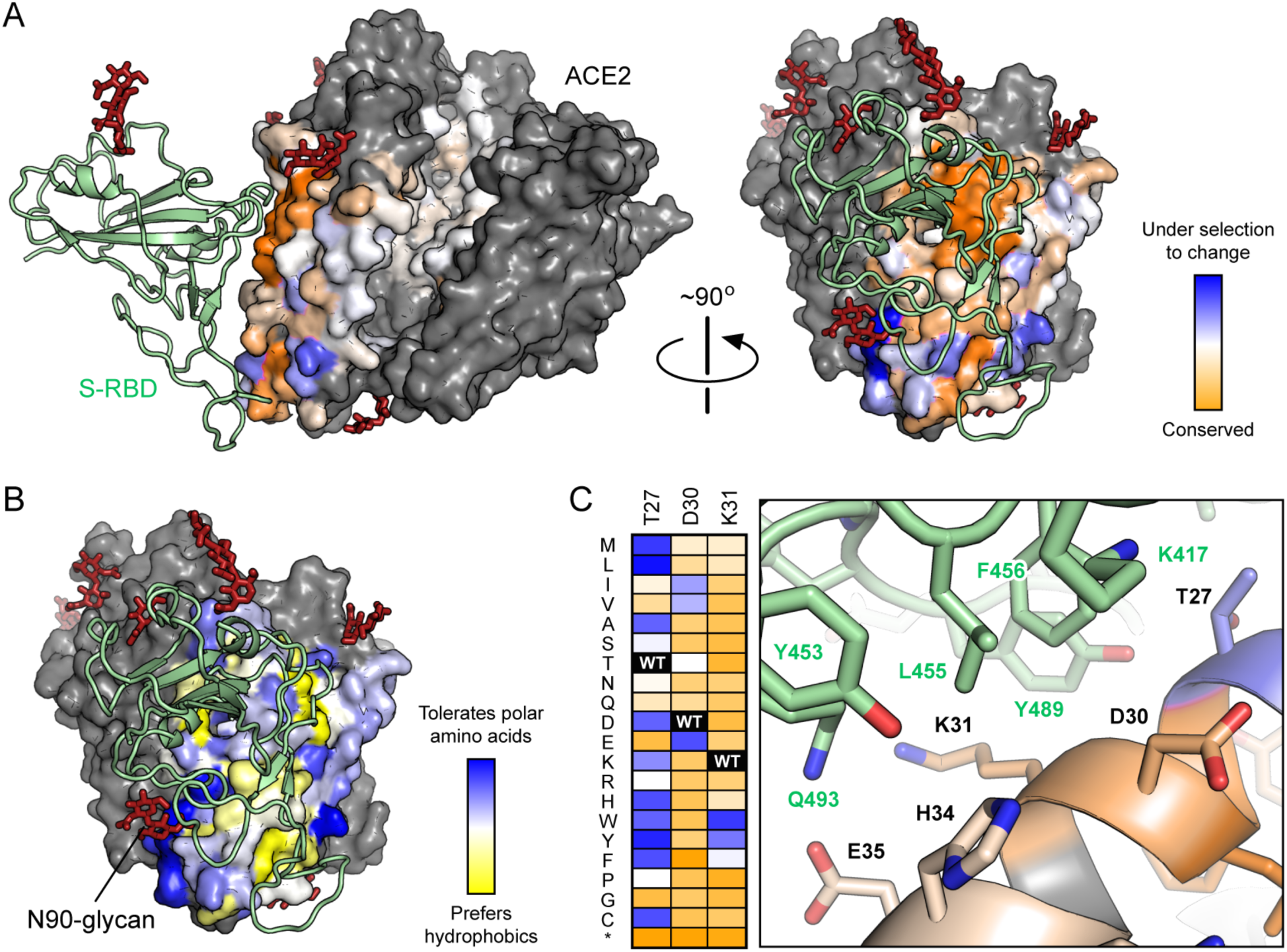
Sequence preferences of ACE2 residues for high binding to the RBD of SARS-CoV-2 S. **(A)** Conservation scores from the nCoV-S-High sorts are mapped to the cryo-EM structure (PDB 6M17) of RBD (pale green ribbon) bound ACE2 (surface). The view at left is looking down the substrate-binding cavity, and only a single protease domain is shown for clarity. Residues conserved for high RBD binding are orange; mutationally tolerant residues are pale colors; residues that are hot spots for enriched mutations are blue; and residues maintained as wild type in the ACE2 library are grey. Glycans are dark red sticks. **(B)** Average hydrophobicity-weighted enrichment ratios are mapped to the RBD-bound ACE2 structure, with residues tolerant of polar substitutions in blue, while residues that prefer hydrophobic amino acids are yellow. **(C)** A magnified view of part of the ACE2 (colored by conservation score as in A) / RBD (pale green) interface. Accompanying heatmap plots log_2_ enrichment ratios from the nCoV-S-High sort for substitutions of ACE2-T27, D30 and K31 from ≤ −3 (depleted) in orange to ≥+3 (enriched) in dark blue.

Two ACE2 residues, N90 and T92 that together form a consensus N-glycosylation motif, are notable hot spots for enriched mutations (Fig. 2 and 4A). Indeed, all substitutions of N90 and T92, with the exception of T92S which maintains the N-glycan, are highly favorable for RBD binding, and the N90-glycan is thus predicted to partially hinder S/ACE2 interaction. This result may be dependent on the chemical nature of glycan moieties attached in different cell types.

Mining the data identifies many ACE2 mutations that are enriched for RBD binding. For instance, there are 122 mutations to 35 positions in the library that have log_2_ enrichment ratios >1.5 in the nCoV-S-High sort. By comparing the mutational landscape with human genetic diversity, it has been proposed that ACE2 polymorphisms are relevant to COVID-19 pathogenesis and transmission (*36*, *37*), although this will require further validation. At least a dozen ACE2 mutations at the structurally characterized interface enhance RBD binding, and will be useful for engineering highly specific and tight protein or peptide binders of SARS-CoV-2 S. The molecular basis for how some of these mutations enhance RBD binding can be rationalized from the RBD-bound cryo-EM structure (Fig. 4C): hydrophobic substitutions of ACE2-T27 increase hydrophobic packing with aromatic residues of S, ACE2-D30E extends an acidic side chain to reach S-K417, and aromatic substitutions of ACE2-K31 contribute to an interfacial cluster of aromatics. A search for affinity-enhancing mutations in ACE2 using targeted mutagenesis recently identified D30E (*38*), providing independent confirmation of the methods used here.

Attention was also drawn to mutations in the second shell and farther from the interface that do not directly contact S but instead have putative structural roles. For example, proline substitutions were enriched at five library positions (S19, L91, T92, T324 and Q325) where they might entropically stabilize the first turns of helices. Proline was also enriched at H34, where it may enforce the central bulge in α1. Multiple mutations were also enriched at buried positions where they will change local packing (e.g. A25V, L29F, W69V, F72Y and L351F). The selection of ACE2 variants for high binding signal therefore not only reports on affinity, but also on presentation at the membrane of folded structure recognized by SARS-CoV-2 S. Whether these mutations selectively stabilize a virus-recognized local structure in ACE2 versus the global protein fold is unclear.

Thirty single substitutions highly enriched in the nCoV-S-High sort were validated by targeted mutagenesis (Fig. 5). Binding of RBD-sfGFP to full length ACE2 mutants increased compared to wild type, yet improvements were small and most apparent on cells expressing low ACE2 levels (Fig. 5A). Differences in ACE2 expression between the mutants also correlated with total levels of bound RBD-sfGFP (Fig. 5C). To rapidly assess mutations in a format more relevant to therapeutic and diagnostic development, the soluble ACE2 protease domain was fused to sfGFP. Expression levels of sACE2-sfGFP were qualitatively evaluated by fluorescence of the transfected cultures (Fig. 6A), and binding of sACE2-sfGFP to full length S expressed at the plasma membrane was measured by flow cytometry (Fig. 6B). A single substitution (T92Q) that eliminates the N90 glycan gave a small increase in binding signal (Fig. 6B). Focusing on the most highly enriched substitutions in the selection for S binding that were also spatially segregated to minimize negative epistasis (*39*), combinations of mutations in sACE2 gave large increases in S binding (Table 1 and Fig. 6B). While this assay only provides relative differences, the combinatorial mutants have enhanced binding by at least an order of magnitude. Unexplored combinations of mutations may have even greater effects.

**Table 1.** Combinatorial mutants of sACE2.

| Variant | Mutations |
| --- | --- |
| sACE2.v1 | H34A, T92Q, Q325P, A386L |
| sACE2.v2 | T27Y, L79T, N330Y, A386L |
| sACE2.v2.1 | L79T, N330Y, A386L |
| sACE2.v2.2 | T27Y, N330Y, A386L |
| sACE2.v2.3 | T27Y, L79T, A386L |
| sACE2.v2.4 | T27Y, L79T, N330Y |
| sACE2.v3 | A25V, T27Y, T92Q, Q325P, A386L |
| sACE2.v4 | H34A, L79T, N330Y, A386L |
| sACE2.v5 | A25V, T92Q, A386L |
| sACE2.v6 | T27Y, Q42L, L79T, T92Q, Q325P, N330Y, A386L |

**Figure 5.**
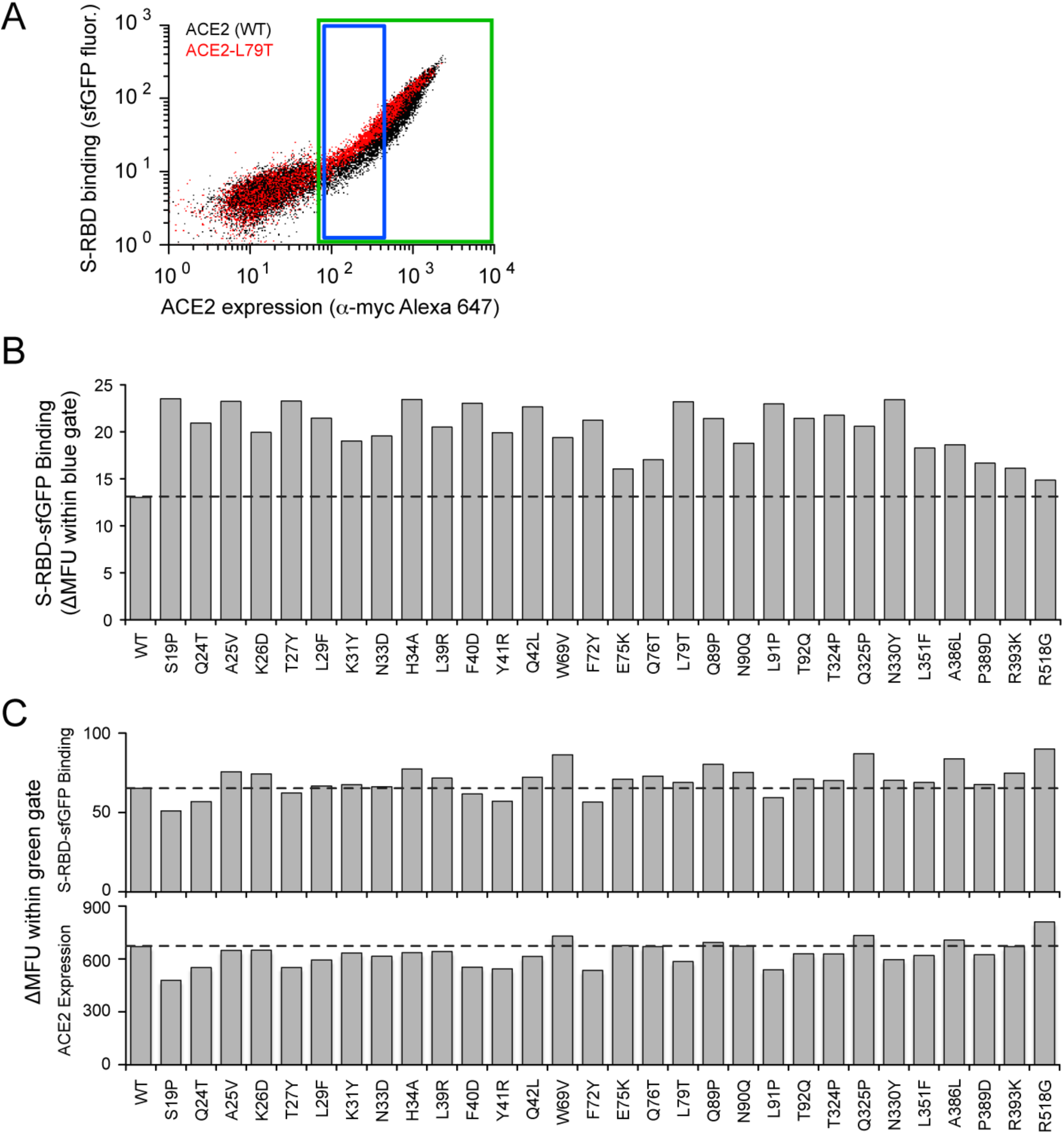
Single amino acid substitutions in ACE2 predicted from the deep mutational scan to increase RBD binding have small effects. **(A)** Expi293F cells expressing full length ACE2 were stained with RBD-sfGFP-containing medium and analyzed by flow cytometry. Data are compared between wild type ACE2 (black) and a single mutant (L79T, red). Increased RBD binding is most discernable in cells expressing low levels of ACE2 (blue gate). In this experiment, ACE2 has an extracellular N-terminal myc tag upstream of residue S19 that is used to detect surface expression. **(B)** RBD-sfGFP binding was measured for 30 single amino acid substitutions in ACE2. Data are GFP mean fluorescence in the low expression gate (blue gate in panel A) with background fluorescence subtracted. **(C)** RBD-sfGFP binding measured for the total ACE2-positive population (green gate in panel A) is shown in the upper graph, while the lower graph plots ACE2 expression measured by detection of the extracellular myc tag. Total RBD-sfGFP binding correlates with total ACE2 expression, and differences in binding between the mutants are therefore most apparent only after controlling for expression levels as in panel A.

**Figure 6.**
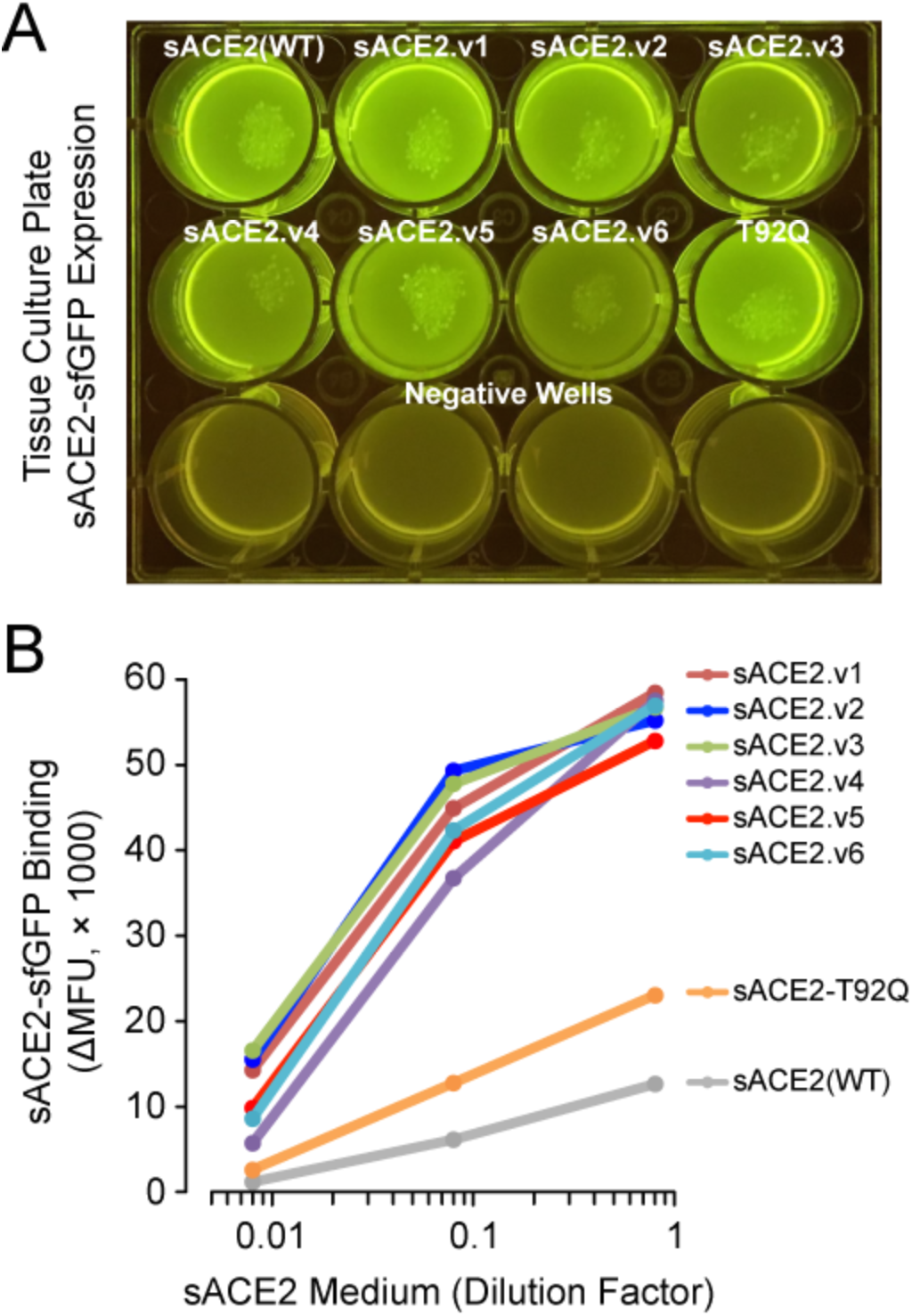
Engineered sACE2 with enhanced binding to S. **(A)** Expression of sACE2-sfGFP mutants was qualitatively evaluated by fluorescence of the transfected cell cultures. **(B)** Cells expressing full length S were stained with dilutions of sACE2-sfGFP-containing media and binding was analyzed by flow cytometry.

A single variant, sACE2.v2, was chosen for purification and further characterization (Fig. 7). This variant was selected because it was well expressed fused to sfGFP and maintains the N90-glycan, and will therefore present a surface that more closely matches native sACE2 to minimize immunogenicity. The yield of sACE2.v2 was lower than the wild type protein when purified as an 8his-tagged protein (20% lower) or as an IgG1-Fc fusion (60% lower), and by analytical size exclusion chromatography (SEC) a small fraction of sACE2.v2 was found to aggregate after incubation at 37 °C for 40 h (Fig. 7D). Otherwise, sACE2.v2 was indistinguishable from wild type by SEC (Fig. 7C).

**Figure 7.**
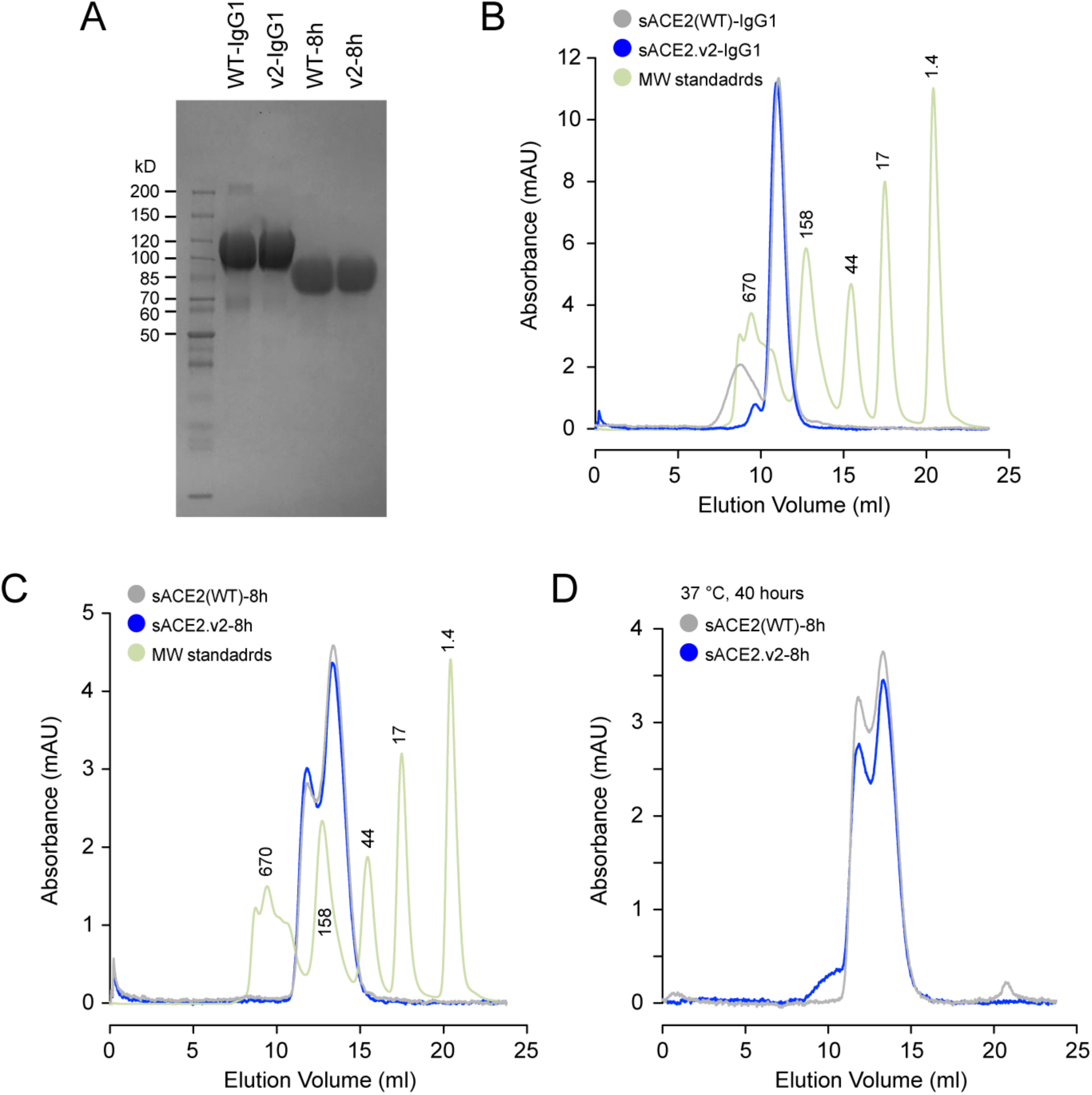
Analytical SEC of purified sACE2 proteins. **(A)** Purified sACE2 proteins (10 μg) were separated on a 4-20% SDS-polyacrylamide gel and stained with coomassie. **(B)** Analytical SEC of IgG1-fused wild type sACE2 (grey) and sACE2.v2 (blue). Molecular weights (MW) of standards (green) are indicated in kD above the peaks. Absorbance of the MW standards is scaled for clarity. **(C)** Analytical SEC of 8his-tagged proteins. The major peak corresponds to the expected MW of a monomer. A dimer peak is also observed, although its abundance differs between independent protein preparations (compare to Figure 10D). **(D)** Soluble ACE2-8h proteins were incubated at 37 °C for 40 h and analyzed by SEC.

In flow cytometry experiments using purified 8his-tagged sACE2, only sACE2.v2-8h was found to bind strongly to full length S at the cell surface, suggestive that wild type sACE2 has a high off-rate that causes dissociation during sample washing (Fig. 8A and 9). Differences between wild type and the variant were less pronounced in the context of an IgG1-Fc fusion (Fig. 8A and 9), indicating that avidity masks gains in binding of the mutant, again suggestive that there are off-rate differences between wild type and variant sACE2. Soluble ACE2.v2-8h outcompetes wild type sACE2-IgG1 for binding to S-expressing cells, yet wild type sACE2-8h does not outcompete sACE2-IgG1 even at 10-fold higher concentrations (Fig. 8B). These results align with a study showing that while wild type sACE2 is highly effective at inhibiting SARS-CoV-2 replication in cell lines and organoids, extremely high concentrations are required (*24*). Cell experiments were supported by biolayer interferometry (BLI), in which IgG1-Fc fused RBD was captured on a biosensor surface and the association and dissociation kinetics of 8his-tagged sACE2 were determined. The K_D_ of wild type sACE2-8h for the RBD was 140 to 150 nM (Fig. 8C and 10F), slightly higher than the K_D_ values reported by others that range from 1 to 50 nM (8, 10)(*40*). Variant sACE2.v2 had 65-fold tighter affinity for the RBD, almost entirely due to a slower off-rate (Fig. 8D). Across all experiments, whether ACE2 is purified as a 8his-tagged protein or used as a sfGFP-fusion in expression medium, and whether full-length S is expressed on the plasma membrane or the isolated RBD is immobilized on a biosensor surface, the characterized sACE2.v2 variant consistently shows one to two orders of magnitude tighter binding. These experiments support the key discovery from deep mutagenesis that mutations in human ACE2 exist that increase binding to S of SARS-CoV-2.

**Figure 8.**
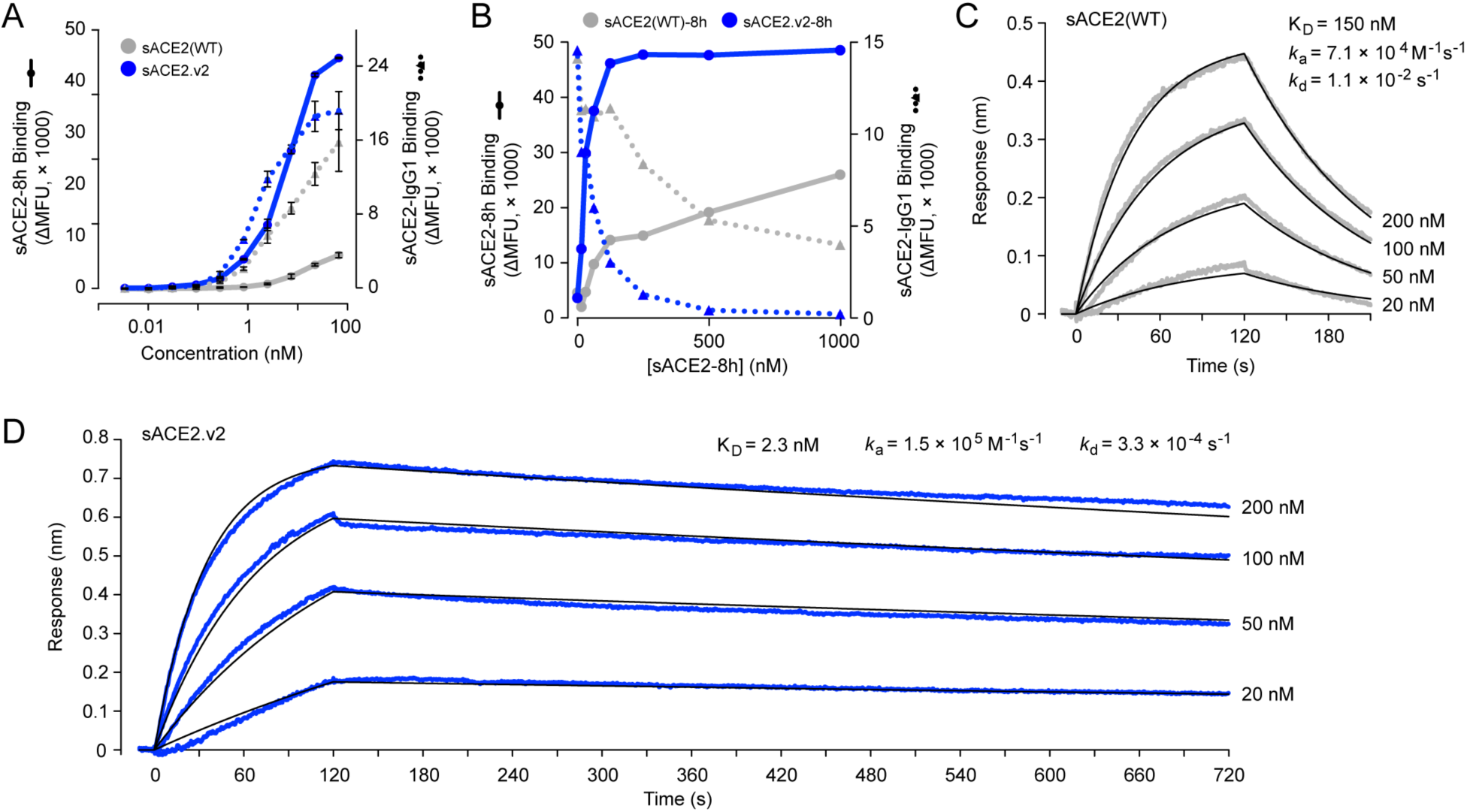
A variant of sACE2 with high affinity for S. **(A)** Expi293F cells expressing full length S were incubated with purified wild type sACE2 (grey) or sACE2.v2 (blue) fused to 8his (solid lines) or IgG1-Fc (broken lines). After washing and staining with secondary antibodies, bound protein was detected by flow cytometry. Data are mean fluorescence units (MFU) of the total cell population after subtraction of background autofluorescence. n = 2, error bars represent range. **(B)** Binding of 100 nM wild type sACE2-IgG1 (broken lines) was competed with wild type sACE2-8h (solid grey line) or sACE2.v2-8h (solid blue line). The competing proteins were added simultaneously to cells expressing full length S, and bound proteins were detected by flow cytometry. **(C)** BLI kinetics of wild type sACE2-8h association (t = 0 to 120 s) and dissociation (t > 120 s) with immobilized RBD-IgG1. Compare to an independent protein preparation in Figure 10F. **(D)** Kinetics of sACE2.v2-8h binding to immobilized RBD-IgG1 measured by BLI.

**Figure 9.**
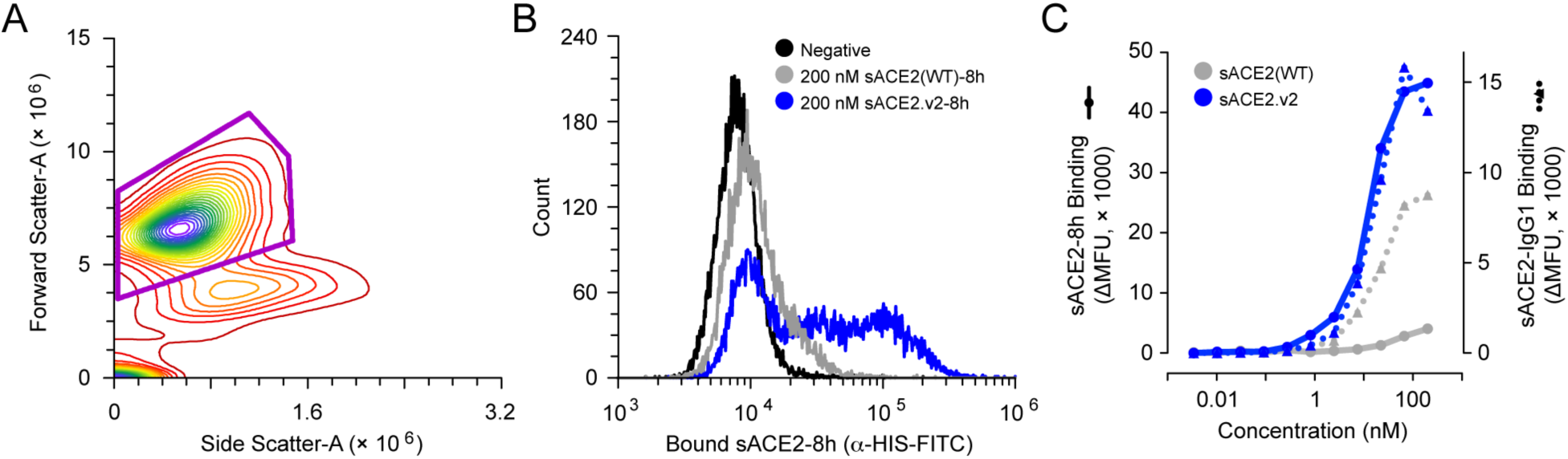
Flow cytometry measurements of sACE2 binding to myc-tagged S expressed at the plasma membrane. **(A)** Expi293F cells expressing full length S, either untagged (Figure 8A) or with an extracellular myc epitope tag (this Figure), were gated by forward-side scattering properties for the main cell population (purple gate). **(B)** Histograms showing representative raw data from flow cytometry analysis of myc-S-expressing cells incubated with 200 nM wild type sACE2-8h (grey) or sACE2.v2 (blue). After washing, bound protein was detected with a fluorescent anti-HIS-FITC secondary. Fluorescence of myc-S-expressing cells treated without sACE2 is black. **(C)** Binding of purified wild type sACE2 (grey) or sACE2.v2 (blue) fused to 8his (solid lines) or IgG1-Fc (broken lines) to cells expressing myc-S.

To address the decreased expression of sACE2.v2, it was hypothesized that the mutational load is too high. Each of the four mutations in sACE2.v2 was reverted back to the wild type identity (Table 1), and binding to full length S at the cell surface was found to remain tight using rapid screening of sACE2-sfGFP-containing expression medium (Fig. 10A). Expression was rescued to varying degrees (Fig. 10B), and one of the variants (sACE2.v2.4 with mutations T27Y, L79T and N330Y) was purified with higher yields than wild type (20% and 80% higher for 8his and IgG1-Fc tagged proteins, respectively), although some protein remained aggregated after storage at 37 °C for 60 h (Figure 10D and 10E). A second reversion variant (sACE2.v2.2 with mutations T27Y, N330Y and A386L) has a more hydrophobic surface and higher propensity to partially aggregate after storage at 37 °C, and therefore the partial storage instability may be intrinsically linked to increased hydrophobicity of the mutated ACE2 surface. By BLI, sACE2.v2.4 displayed tight nanomolar binding to the RBD (Fig. 10G), demonstrating that high affinity can be achieved without compromising protein expression and purification. Further sequence optimization focused on whether to include the ACE2 neck domain for stable dimer formation is warranted.

**Figure 10.**
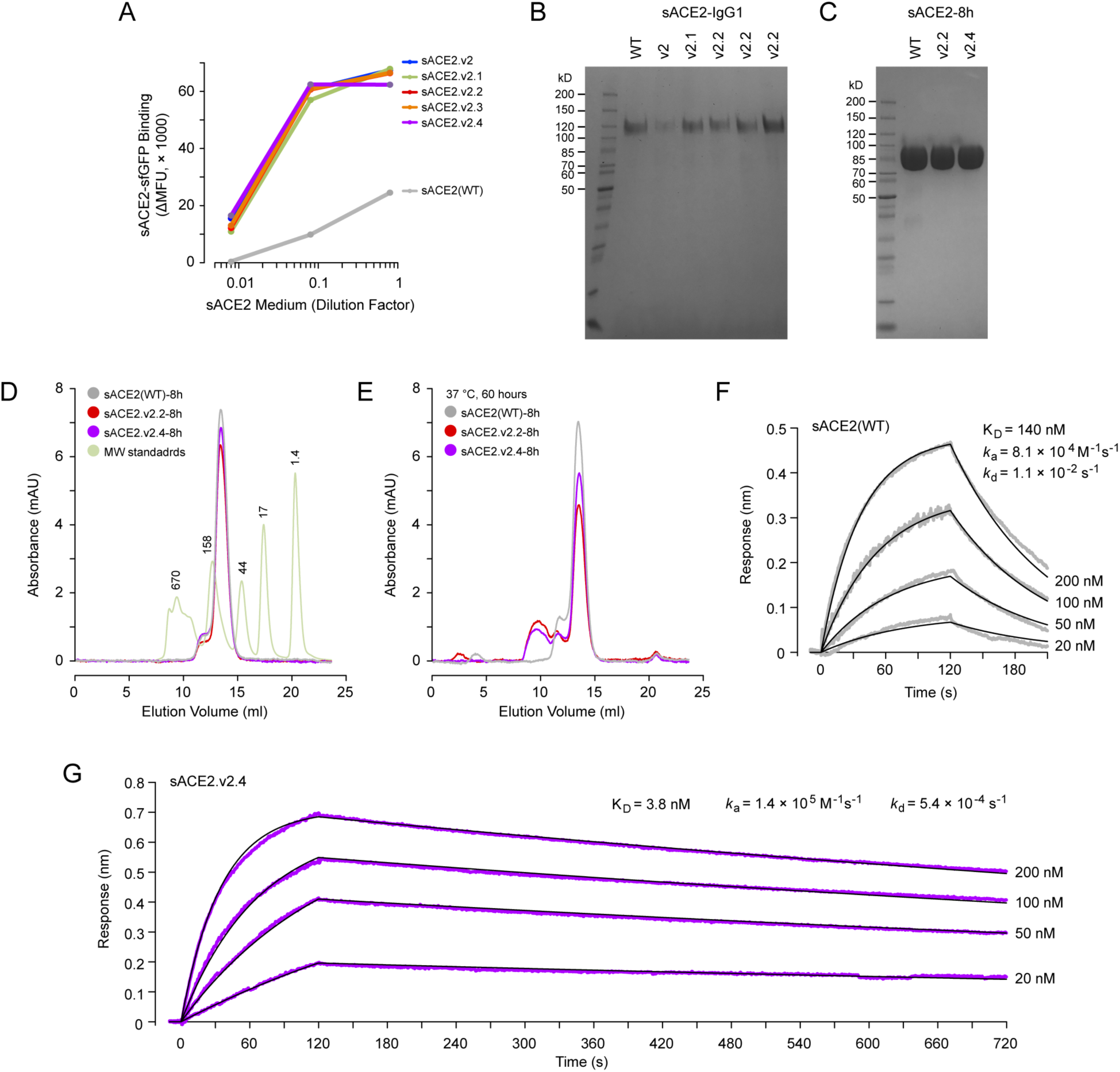
Optimization of a high affinity sACE2 variant for improved yield. **(A)** Dilutions of sACE2-sfGFP-containing media were incubated with Expi293F cells expressing full length S. After washing, bound sACE2-sfGFP was analyzed by flow cytometry. **(B)** Coomassie-stained SDS-polyacrylamide gel compares the yield of sACE2-IgG1 variants purified from expression medium by protein A resin. **(C)** Coomassie-stained gel of purified sACE2-8h variants (10 μg per lane). **(D)** By analytical SEC, sACE2.v2.2-8h (red) and sACE2.v2.4-8h (purple) are indistinguishable from wild type sACE2-8h (grey). The absorbance of MW standards (green) is scaled for clarity, with MW indicated above the elution peaks in kD. **(E)** Analytical SEC after storage of the proteins at 37 °C for 60 h. **(F)** Wild type sACE2-8h association (t = 0 to 120 s) and dissociation (t > 120 s) with immobilized RBD-IgG1 measured by BLI. Data are comparable to a second independent preparation of sACE2-8h shown in Figure 8C. **(G)** BLI kinetics of sACE2.v2.4-8h with immobilized RBD-IgG1.

While deep mutagenesis of viral proteins in replicating viruses has been extensively pursued to understand escape mechanisms from drugs and antibodies, the work here shows how deep mutagenesis can be directly applicable to anti-viral protein design when the selection method is decoupled from virus replication and focused on host factors.

## METHODS

### Plasmids

The mature polypeptide (a.a. 19-805) of human ACE2 (GenBank NM_021804.1) was cloned in to the NheI-XhoI sites of pCEP4 (Invitrogen) with a N-terminal HA leader (MKTIIALSYIFCLVFA), myc-tag, and linker (GSPGGA). Soluble ACE2 fused to superfolder GFP (*34*) was constructed by genetically joining the protease domain (a.a. 1-615) of ACE2 to sfGFP (GenBank ASL68970) via a gly/ser-rich linker (GSGGSGSGG), and pasting between the NheI-XhoI sites of pcDNA3.1(+) (Invitrogen). Equivalent sACE2 constructs were cloned with a GSG linker and 8 histidine tag, or a GS linker and the Fc region of IgG1 (a.a. D221-K447). A synthetic human codon-optimized gene fragment (Integrated DNA Technologies) for the RBD (a.a. 333-529) of SARS-CoV-2 S (GenBank YP_009724390.1) was N-terminally fused to a HA leader and C-terminally fused to either superfolder GFP, the Fc region of IgG1 or a 8 histidine tag. Assembled DNA fragments were ligated in to the NheI-XhoI sites of pcDNA3.1(+). Human codon-optimized full length S was subcloned from pUC57-2019-nCoV-S(Human) (Molecular Cloud), both untagged (a.a. 1-1273) and with a N-terminal HA leader (MKTIIALSYIFCLVFA), myc-tag and linker (GSPGGA) upstream of the mature polypeptide (a.a. 16-1273).

### Tissue Culture

Expi293F cells (ThermoFisher) were cultured in Expi293 Expression Medium (ThermoFisher) at 125 rpm, 8 % CO_2_, 37 °C. For production of RBD-sfGFP, RBD-IgG1, sACE2-8h and sACE2-IgG1, cells were prepared to 2 × 10^6^ / ml. Per ml of culture, 500 ng of plasmid and 3 ¼g of polyethylenimine (MW 25,000; Polysciences) were mixed in 100 ¼ of OptiMEM (Gibco), incubated for 20 minutes at room temperature, and added to cells. Transfection Enhancers (ThermoFisher) were added 18-23 h post-transfection, and cells were cultured for 4-5 days. Cells were removed by centrifugation at 800 × g for 5 minutes and medium was stored at −20 °C. After thawing and immediately prior to use, remaining cell debris and precipitates were removed by centrifugation at 20,000 × g for 20 minutes. Plasmids for expression of sACE2-sfGFP protein were transfected in to Expi293F cells using Expifectamine (ThermoFisher) according to the manufacturer’s directions, with Transfection Enhancers added 22-^1^/_2_ h post-transfection, and medium supernatant harvested after 60 h.

### Deep mutagenesis

117 residues within the protease domain of ACE2 were diversified by overlap extension PCR (*41*) using primers with degenerate NNK codons. The plasmid library was transfected in to Expi293F cells using Expifectamine under conditions previously shown to typically give no more than a single coding variant per cell (*32*, *33*); 1 ng coding plasmid was diluted with 1,500 ng pCEP4-ΔCMV carrier plasmid per ml of cell culture at 2 × 10^6^ / ml, and the medium was replaced 2 h post-transfection. The cells were collected after 24 h, washed with ice-cold PBS supplemented with 0.2 % bovine serum albumin (PBS-BSA), and incubated for 30 minutes on ice with a 1/20 (replicate 1) or 1/40 (replicate 2) dilution of medium containing RBD-sfGFP into PBS-BSA. Cells were co-stained with anti-myc Alexa 647 (clone 9B11, 1/250 dilution; Cell Signaling Technology). Cells were washed twice with PBS-BSA, and sorted on a BD FACS Aria II at the Roy J. Carver Biotechnology Center. The main cell population was gated by forward/side scattering to remove debris and doublets, and DAPI was added to the sample to exclude dead cells. Of the myc-positive (Alexa 647) population, the top 67% were gated (Fig. 1B). Of these, the 15 % of cells with the highest and 20% of cells with the lowest GFP fluorescence were collected (Fig. 1D) in tubes coated overnight with fetal bovine serum and containing Expi293 Expression Medium. Total RNA was extracted from the collected cells using a GeneJET RNA purification kit (Thermo Scientific), and cDNA was reverse transcribed with high fidelity Accuscript (Agilent) primed with gene-specific oligonucleotides. Diversified regions of ACE2 were PCR amplified as 5 fragments. Flanking sequences on the primers added adapters to the ends of the products for annealing to Illumina sequencing primers, unique barcoding, and for binding the flow cell. Amplicons were sequenced on an Illumina NovaSeq 6000 using a 2×250 nt paired end protocol. Data were analyzed using Enrich (*42*), and commands are provided in the GEO deposit. Briefly, the frequencies of ACE2 variants in the transcripts of the sorted populations were compared to their frequencies in the naive plasmid library to calculate a log_2_ enrichment ratio and then normalized by the same calculation for wild type. Wild type sequences were neither substantially enriched or depleted, and had log_2_ enrichment ratios of −0.2 to +0.2.

### Flow Cytometry Analysis of ACE2-S Binding

Expi293F cells were transfected with pcDNA3-myc-ACE2, pcDNA3-myc-S or pcDNA3-S plasmids (500 ng DNA per ml of culture at 2 × 10^6^ / ml) using Expifectamine (ThermoFisher). Cells were analyzed by flow cytometry 24 h post-transfection. To analyze binding of RBD-sfGFP to full length myc-ACE2, cells were washed with ice-cold PBS-BSA, and incubated for 30 minutes on ice with a 1/30 dilution of medium containing RBD-sfGFP and a 1/240 dilution of anti-myc Alexa 647 (clone 9B11, Cell Signaling Technology). Cells were washed twice with PBS-BSA and analyzed on a BD LSR II. To analyze binding of sACE2-sfGFP to full length myc-S, cells were washed with PBS-BSA, and incubated for 30 minutes on ice with a serial dilution of medium containing sACE2-sfGFP and a 1/240 dilution of anti-myc Alexa 647 (clone 9B11, Cell Signaling Technology). Cells were washed twice with PBS-BSA and analyzed on a BD Accuri C6, with the entire Alexa 647-positive population gated for analysis. To measure binding of sACE2-IgG1 or sACE2-8h, myc-S or S transfected cells were washed with PBS-BSA and incubated for 30 minutes with the indicated concentrations of purified sACE2 in PBS-BSA. Cells were washed twice, incubated with secondary antibody (1/100 dilution of chicken anti-HIS-FITC polyclonal from Immunology Consultants Laboratory; or 1/250 anti-human IgG-APC clone HP6017 from BioLegend) for 30 minutes on ice, washed twice again, and fluorescence of the total population after gating by FSC-SSC to exclude debris was measured on a BD Accuri C6. Data were processed with FCS Express (De Novo Software) or BD Accuri C6 Software.

### Purification of IgG1-Fc fused proteins

Cleared expression medium was incubated with KANEKA KanCapA 3G Affinity sorbent (Pall; equilibrated in PBS) for 90 minutes at 4 °C. The resin was collected on a chromatography column, washed with 12 column volumes (CV) PBS, and protein eluted with 5 CV 60 mM Acetate pH 3.7. The eluate was immediately neutralized with 1 CV of 1 M Tris pH 9.0, and concentrated with a 100 kD MWCO centrifugal device (Sartorius). Protein was separated on a Superdex 200 Increase 10/300 GL column (GE Healthcare Life Sciences) with PBS as the running buffer. Peak fractions were pooled, concentrated to ~10 mg/ml with excellent solubility, and stored at −80 °C after snap freezing in liquid nitrogen. Protein concentrations were determined by absorbance at 280 nm using calculated extinction coefficients for monomeric, mature polypeptide sequences.

### Purification of 8his-tagged proteins

HisPur Ni-NTA resin (Thermo Scientific) equilibrated in PBS was incubated with cleared expression medium for 90 minutes at 4 °C. The resin was collected on a chromatography column, washed with 12 column volumes (CV) PBS, and protein eluted with a step elution of PBS supplemented with 20 mM, 50 mM and 250 mM imidazole pH 8 (6 CV of each fraction). The 50 mM and 250 mM imidazole fractions were concentrated with a 30 kD MWCO centrifugal device (MilliporeSigma). Protein was separated on a Superdex 200 Increase 10/300 GL column (GE Healthcare Life Sciences) with PBS as the running buffer. Peak fractions were pooled, concentrated to ~5 mg/ml with excellent solubility, and stored at −80 °C after snap freezing in liquid nitrogen.

### Analytical SEC

Proteins (200 μl at 2 μM) were separated on a Superdex 200 Increase 10/300 GL column (GE Healthcare Life Sciences) equilibrated in PBS. MW standards were from Bio-Rad.

### Biolayer Interferometry

Hydrated anti-human IgG Fc biosensors (Molecular Devices) were dipped in expression medium containing RBD-IgG1 for 60 s. Biosensors with captured RBD were washed in assay buffer, dipped in the indicated concentrations of sACE2-8h protein, and returned to assay buffer to measure dissociation. Data were collected on a BLItz instrument and analyzed with a 1:1 binding model using BLItz Pro Data Analysis Software (Molecular Devices). The assay buffer was 10 mM HEPES pH 7.6, 150 mM NaCl, 3 mM EDTA, 0.05% polysorbate 20, 0.5% non-fat dry milk (Bio-Rad).

### Reagent and data availability

Plasmids are deposited with Addgene under IDs 141183-5, 145145-78, 149268-71 and 149663-8. Raw and processed deep sequencing data are deposited in NCBI’s Gene Expression Omnibus (GEO) with series accession no. GSE147194.

## Supporting information

Processed deep mutagenesis data.

## ACKNOWLEDGEMENTS

Staff at the UIUC Roy J. Carver Biotechnology Center assisted with FACS and Illumina sequencing. Hannah Choi and Krishna Narayanan (University of Illinois) assisted with plasmid preparation. Kui Chan (Orthogonal Biologics Inc) helped characterize sACE2.v2.4. The development of deep mutagenesis to study virus-receptor interactions was supported by NIH award R01AI129719.

